# Multi-omics co-localization with genome-wide association studies reveals a context-specific genetic mechanism at a childhood onset asthma risk locus

**DOI:** 10.1101/593558

**Authors:** Marcus M. Soliai, Atsushi Kato, Catherine T. Stanhope, James E. Norton, Katherine A. Naughton, Aiko I. Klinger, Robert C. Kern, Bruce K. Tan, Robert P. Schleimer, Dan L. Nicolae, Jayant M. Pinto, Carole Ober

## Abstract

**Background:** Genome-wide association studies (GWASs) have identified thousands of variants associated with asthma and other complex diseases. However, the functional effects of most of these variants are unknown. Moreover, GWASs do not provide context-specific information on cell types or environmental factors that affect specific disease risks and outcomes. To address these limitations, we used an upper airway (sinonasal) epithelial cell culture model to assess transcriptional and epigenetic responses to an asthma-promoting pathogen, rhinovirus (RV), and provide context-specific functional annotations to variants discovered in GWASs of asthma.

**Methods:** Using genome-wide genetic, gene expression and DNA methylation data in vehicle- and RV-treated airway epithelial cells (AECs) from 104 individuals, we mapped *cis* expression and methylation quantitative trait loci (*cis*-eQTLs and *cis*-meQTLs, respectively) in each condition. A Bayesian test for co-localization between AEC molecular QTLs and adult onset and childhood onset GWAS variants was used to assign function to variants associated with asthma. Mendelian randomization was applied to demonstrate DNA methylation effects on gene expression at asthma colocalized loci.

**Results:** Co-localization analyses of airway epithelial cell molecular QTLs with asthma GWAS variants revealed potential molecular disease mechanisms of asthma, including QTLs at the *TSLP* locus that were common to both exposure conditions and to both childhood and adult onset asthma, as well as QTLs at the 17q12-21 asthma locus that were specific to RV exposure and childhood onset asthma, consistent with clinical and epidemiological studies of these loci.

**Conclusion:** This study provides information on functional effects of asthma risk variants in airway epithelial cells and insight into a disease-relevant viral exposure that modulates genetic effects on transcriptional and epigenetic responses in cells and on risk for asthma in GWASs.

## Background

Over the past decade, genome-wide association studies (GWASs) have identified over 60 asthma susceptibility loci at genome-wide levels of significance (p<5×10^−8^) [1], with a locus at 17q12-21 being the most replicated and most significant asthma susceptibility locus in childhood onset asthma (reviewed in [2]). Although asthma is typically diagnosed based on clinical symptoms, such as wheeze, cough, and shortness of breath, it is actually comprised of many overlapping phenotypes and distinct endotypes with shared as well as unique genetic and environmental risk factors. For example, individuals with asthma differ with respect to age of onset, environmental triggers of exacerbations, response to medications, obesity, and co-occurrence with allergic diseases and other conditions. Recently, Pividori et al. reported 61 independent asthma loci, 23 of which were specific to childhood onset asthma, one that was specific to adult onset asthma, and 37 that were associated with risk for both childhood onset and adult onset asthma [1]. Gene and tissue enrichment patterns at these risk loci suggested that epithelial cells (skin) and lung as primary etiological drivers of childhood onset and adult onset asthma, respectively, while blood (immune) cell gene expression enrichments were shared by both. However, GWASs do not generally consider tissue-or other environment-specific effects, or gene by environment interactions. Moreover, epigenetic patterning may mediate the effects of exposures on gene expression and disease risk, yet such studies have only rarely been integrated with GWAS of asthma [3].

A challenge in interpreting GWAS results is that over 90% of disease-associated variants are located in non-protein-coding regions of the genome [4], which are enriched for chromatin signatures suggestive of enhancers [4] and for expression quantitative trait loci (eQTLs) [4–6]. SNPs associated in GWASs that also have functional annotations are more likely to be causal variants, underlying disease pathophysiology through their effects on gene regulation. However, identifying causal variants and their target genes at associated loci has been challenging, and the functions of most SNPs associated with diseases in GWASs remain unknown. Databases such as GTEx, ENCODE, and ROADMAP have been used to annotate GWAS SNPs and predict molecular mechanisms through which risk variants affect disease phenotypes [3, 6–8]. But while these resources have provided important insights into the interpretation of GWAS results, they do not include all cell types relevant to all diseases or information on environmental exposures that influence disease outcomes. As a result, annotations of asthma GWAS variants have been largely limited to studies in transformed B cells lines, blood (immune) cells, and whole lung tissue [1, 9, 10].

*In vitro* cell models provide an opportunity to address these limitations by characterizing genetic and molecular responses to environmental exposures in cells from disease-relevant tissues, and identifying genotypes that modify these responses [11, 12]. Joint analysis of datasets (e.g. eQTLs and GWASs) can identify variants associated with both disease risk and molecular traits as candidate causal variants that contribute to mechanisms of disease pathophysiology. A multi-trait co-localization method (*moloc*) [13] was developed to integrate summary data from GWAS and multiple molecular QTL datasets and identify candidate regulatory drivers of complex phenotypes.

Here, we report the results of a multi-omics co-localization study to identify condition-specific regulatory effects of asthma risk variants using an epithelial cell model of viral response. Because airway epithelium forms a barrier to inhaled exposures, we used an *in vitro* upper airway (sinonasal) epithelial cell model of transcriptional and epigenetic responses to rhinovirus (RV). Primary infection of RV occurs in the nasal epithelium, and RV is a major contributor to asthma inception in young children [14] and asthma exacerbations throughout life [15, 16], underscoring its importance as a contextual promoter of asthma pathophysiology. We demonstrate a specific enrichment of childhood onset asthma GWAS SNPs among airway epithelial molecular QTLs, consistent with the important role that the epithelial barrier plays in the inception of asthma in childhood [1, 17, 18]. Our integrative multi-omics approach suggests an environment-specific mechanism of asthma pathogenesis at the 17q12-21 asthma locus in childhood onset asthma, and a molecular mechanism shared between childhood onset and adult onset GWASs in the *TSLP* gene at chromosome 5q22, highlighting complementary roles of the airway epithelium in the pathogenesis of asthma.

## Methods

### Ethics statement

Study participants were recruited between March 2012 and August 2015, and nasal specimens were collected as part of routine endoscopic sinonasal surgeries at Northwestern University Feinberg School of Medicine. Informed written consent was obtained from each study participant and randomly generated ID codes were assigned to all samples thereby preserving the participant’s anonymity and privacy. This study was approved by the institutional review boards at Northwestern University Feinberg School of Medicine and the University of Chicago.

### Sample collection and composition

Sinonasal epithelial cells were obtained by brushing the uncinate process collected at elective surgery at Northwestern University from 56 males, 39 females, ages 18 – 73 years old (mean age 44), self-reported ethnicities as Caucasian (64%), Black (17%), Hispanic (13%), and “other” (6%), and 43 asthmatics (current or prior status) and 61 non-asthmatics. Blood samples for genotyping were collected from study participants. A summary of the study design is shown in Fig. S1.

### Upper airway epithelial cell culture and RV treatment

After isolation, nasal airway epithelial cells were cultured in bronchial epithelial cell growth medium (Lonza, BEGM BulletKit, catalog number CC-3170) to near confluence, then frozen at −80°C and stored in Liquid Nitrogen. Cells were subsequently thawed and cultured in collagen-coated (PureCol, INAMED BioMaterials, catalog number 5,409, 3 mg/mL, 1:15 dilution) tissue culture plates (6 wells of 2× 12 well plates) using BEGM overnight at 37°C and 5% CO_2_. In preparation for rhinovirus (HRV-16; RV) infection/stimulation, plates at 50-60% confluency were incubated overnight in BEGM without hydrocortisone (HC) followed by a two-hour RV infection at a multiplicity of infection (MOI) of 2 and vehicle treatment (Bronchial epithelial cell basal medium (BEBM) + Gentamicin/Amphotericin) at 33°C (low speed rocking, ~15 RPM). RV- and vehicle-treated cells were washed and then were cultured at 33°C for 46 hours (48 hours total) in BEGM without HC.

### Genotyping and imputation

DNA was extracted from whole blood or sinus tissue (if no blood was available) with the Macherey-Nagel NucleoSpin Blood L or NucleoSpin Tissue L Extraction kits, respectively, and quantified with the NanoDrop ND1000. Genotyping of all study participants was performed using the Illumina Infinium HumanCore Exome+Custom Array (550,224 SNPs). After quality control (QC) (excluding SNPs with HWE < 0.0001 by race/ethnicity, call rate < 0.95, MAF < 0.05 and individuals with genotype call rates < 0.05), 529,993 markers for 104 individuals were available for analysis. Ancestry principal component analysis (PCA) was performed using 676 ancestry informative markers included on the array that overlap with the HapMap release 3 (Fig. S2).

Phasing and imputation were performed using the ShapeIt2 [19] and Impute2 [20] software packages, respectively. Variants were imputed in 5 Mb windows across the genome against the 1000 Genomes Phase 3 haplotypes (Build 37; October 2014). Individuals were categorized into two groups based on the k-means clustering of ancestry PCs, using the kmeans() function in R; individuals were grouped as European or African American based on how they related to the HapMap reference panel and means clustering of their ancestry PCs (Fig. S2). After imputation, both groups were merged and QC was performed with gtool [21]. X and Y chromosome-linked SNPs and SNPs that did not meet the QC criteria (info score < 0.8, MAF < 0.05, missingness > 0.05 and a probability score < 0.9) were excluded from analyses. Probability scores were converted to dosages for 6,665,552 of the remaining sites used in downstream analyses.

### RNA extraction and sequencing

Following RV and vehicle treatments, RNA from cells underwent extraction and purification using the QIAGEN AllPrep DNA/RNA Kit. RNA quality and quantity were measured at the University of Chicago Functional Genomics Core using the Agilent RNA 6000 Pico assay and the Agilent 2100 Bioanalyzer. RNA integrity numbers (RIN) were greater than 7.7 for all samples. cDNA libraries were constructed using the Illumina TruSeq RNA Library Prep Kit v2 and sequenced on the Illumina HiSeq 2500 System (50 bp, single-end); RNA sequencing was completed at the University of Chicago Genomics Core. Subsequently, we checked for potential sample contamination and sample swaps using the publicly available software VerifyBamID (http://genome.sph.umich.edu/wiki/VerifyBamID) [22] for cells from all 104 individuals included in each treatment condition. We did not detect any cross-contamination between samples but we did identify one sample swap between individuals, which we subsequently corrected.

Sequences were mapped to the human reference genome (hg19) and reads per gene were quantified using the Spliced Transcripts Alignment to a Reference (STAR) [23] software. X,Y, and mitochondrial chromosome genes, and low count data (genes < 1CPM) were removed prior to normalization via the trimmed mean of M-values method (TMM) and variance modeling (voom) [24]; samples contain > 8M mapped reads. Principle components analysis (PCA) identified biological and technical sources of variation in the voom-normalized RNA-seq reads. We identified contributors to batch and other technical effects (days in liquid nitrogen, experimental culture days, cell culture batches, RNA concentration, RNA fragment length, technician, sequencing pool). Additionally, unknown sources of variation were predicted with the Surrogate Variable Analysis (SVA) [25] package in R where 15 surrogate variables (SVs) were estimated for the samples that were included in the experiment. Voom-normalized RNA-seq data were then adjusted for technical effects, SVs, sex, and ancestry PCs (1-3) using the function removeBatchEffect() from the R package limma [26]. Treatment responses in epithelial cells were detected in the combined sample with 6,650 differentially expressed genes identified at a FDR≤0.01 (Fig. S3).

### DNA extraction and methylation profiling

Following RV and vehicle treatments, DNA was extracted from cells as described above. DNA methylation profiles for cells from each treatment were measured on the Illumina Infinium MethylationEPIC BeadChip at the University of Chicago Functional Genomics Core. Methylation data were preprocessed using the minfi package [27]. Probes located on sex chromosomes and with detection p-values greater than 0.01 in more than 10% of samples were removed from the analysis; samples with more than 5% missing probes were also removed. A preprocessing control normalization function was applied to correct for raw probe values or background and a Subset-quantile Within Array Normalization (SWAN) [28] was used to correct for technical differences between the Infinium type I and type II probes. Additionally, we removed cross-reactive probes and probes within two nucleotides of a SNP with an MAF greater than 0.05 using the function rmSNPandCH() from the R package DMRcate [29].

PCA identified technical and biological sources of variation in the normalized DNA methylation datasets. We identified contributors to batch and technical effects including array, and cell harvest date. Sex, age, and smoking were significant variables in the PCA. Unknown sources of variation were predicted with the SVA package where we estimated 37 SVs. SWAN and quantile-normalized M-values were then adjusted for batch and technical effects, SVs, sex, age, and smoking using the function removeBatchEffect() in R. Treatment effects were detected in the combined sample with 1,710 differentially methylated CpGs at a FDR<0.10 (Fig. S4).

### eQTL and meQTL analyses

Prior to e/meQTL analysis, voom-transformed gene expression values and normalized methylation M-values were adjusted for technical (array, cell harvest date), biological variables (sex, age, ancestry PCs), as well as smoking and surrogate variables as described above. Linear regression between the permuted genotypes (MAF>0.05) and molecular phenotypes (gene expression and methylation residuals) from each treatment condition was performed with the FastQTL [30] software package within *cis*-window sizes of 1 Mb and 10 kb for eQTL and meQTL analyses, respectively. Nominal passes were conducted for each eQTL and meQTL analysis within FastQTL, and an FDR threshold of 0.10 was applied to adjust for multiple testing within each experimental dataset with the p.adjust() function in R.

A conditional analysis was performed with the QTLtools [31] software package to identify molecular QTLs with independent effects on gene expression and DNA methylation. This was accomplished in two-steps. First, a permutation analysis was performed within a *cis*-window sizes of 1 Mb and 10 kb for eQTL and meQTL analyses, respectively, to derive nominal p-value thresholds per molecular phenotype. Second, a forward-backward stepwise regression is applied to ultimately assign significant variants to independent signals.

### Multivariate adaptive shrinkage analysis (mash)

An Empirical Bayes method of multivariate adaptive shrinkage was applied separately to the eQTL and meQTL data sets as implemented in the R statistical package, mashr (https://github.com/stephenslab/mashr) [32], to produce improved estimates of QTL effects and corresponding significance values in each treatment condition. Mashr implements this in two general steps: 1) identification of pattern sharing, sparsity, and correlation among QTL effects, and 2) integration of these learned patterns to produce improved effects estimates and measures of significance for eQTLs or meQTLs in each treatment condition. To fit the mash model, we first estimated the correlation structure in the null test from a random dataset in which 235,851 and 3,959,482 phenotype-SNP pairs were chosen for eQTLs and meQTLs, respectively, from the FastQTL nominal pass; because mashr is computationally intensive, the number of randomly chosen gene/CpG-SNP pairs were determined based on R’s memory capabilities. The data-driven covariances were then estimated using the ‘top’ mQTL in each gene or CpG results from FastQTL. Posterior summaries were then computed for the ‘top’ eQTL and meQTL results (see [32]). The instructions found in the *mashr* eQTL analysis outline vignette were followed to run mash.

### Enrichment analysis

The R package, GWAS analysis of regulatory or functional information enrichment with LD correction (GARFIELD) [33], was used to quantify enrichment and assess significance of GWAS SNPs among eQTLs and meQTLs. GARFIELD leverages GWAS results with molecular data to identify features relevant to a phenotype of interest, while accounting for LD and matching for genotyped variants, by applying a logistic regression method to derive statistical significance for enrichment. For this study, molecular QTLs were tested for GWAS variant enrichment, estimated as odds ratios and enrichment P-values derived at four GWAS P-value thresholds: 10^−5^, 10^−6^, 10^−7^, and 10^−8^. To demonstrate disease-specificity of our results, we selected summary statistics from four GWASs performed in UK Biobank subjects (Alzheimer’s disease [34], atrial fibrillation [35], height [36], neuroticism [37], in addition to one each for adult onset and childhood onset asthma [1]). Summary statistics from these six GWASs were used for enrichment analyses of the 755,441 molecular QTLs combined from each treatment condition. These non-asthma GWASs were chosen based on similar population backgrounds (European), availability of summary statistics (as of 05/18), and not known or expected to have overlapping genetics with asthma (i.e., excluding diseases with known allergic or autoimmune etiologies).

To assess tissue-specificity of our results, we examined eQTLs from the adrenal gland, frontal cortex, hypothalamus, ovary, and testis from the GTEx database version 7 (http://gtexportal.org) [6], and tested for enrichment of adult onset and childhood onset asthma GWAS SNPs among the epithelial eQTLs from our study combined across treatment conditions. GTEx data were matched with respect to sample size and number of eQTLs with those of the epithelium, with the exception of testis, which was included to show the consistency of the enrichment results despite it being an outlier in regards to both sample size, which was smaller, and number of eQTLs, which was larger. An FDR threshold of 5% and 10% was applied to eQTLs from GTEx and from our study, respectively, for a balanced, unbiased assessment of enrichment. An OR > 1 and a Benjamini-Hochberg (BH) corrected p-value threshold of < 0.05 was used as the significance threshold for enrichment; BH adjusted p-values were calculated using the p.adjust() function in R where ‘n’ was determined by the number of tests in each respective enrichment analysis.

### Co-localization analysis

To estimate the posterior probability association (PPA) that a SNP contributed to the association signal in the GWAS as well as to the eQTL and/or meQTL, we applied a Bayesian statistical framework implemented in the R package multiple-trait-coloc (moloc) [13]. Summary data from adult onset and childhood onset asthma GWASs from [1], along with eQTL and meQTL summary data from cells within each treatment condition (described above), were included in the *moloc* analysis. Each co-localization analysis included summary data from a GWAS and epithelial cell eQTLs and meQTLs from corresponding treatment conditions. Because a genome-wide co-localization analysis was computationally untenable, genomic regions for co-localization were defined using GARFIELD. First, we analyzed the enrichment pattern of e/meSNPs from each treatment condition in adult onset and childhood onset GWASs using the default package settings. Second, we extracted variants driving the enrichment signals at a GWAS p-value threshold of 1×10^−4^. Regions were defined as 2 Mb windows centered around these variants. Only regions with at least 10 SNPs in common between all three datasets or ‘traits’ (GWAS, eQTL, and meQTL) were assessed by moloc and 15 ‘configurations’ of possible variant sharing was computed across these three traits (see [13] for more details). PPAs ≥ 0.70 were considered as evidence for co-localization. Prior probabilities of 1×10^−4^, 1×10^−6^, and 1×10^−7^ were chosen for the association of one, two, or three traits, respectively, as recommended by the authors of moloc.

### Mendelian randomization

Mendelian Randomization was performed using the ivreg2 function in R (https://www.r-loggers.com/an-ivreg2-function-for-r/) which applies a 2-stage least squares regression (2SLS), as implemented in [38]. We used the only co-localized triplet (eQTL-meQTL-GWAS) to assess the causal effects of DNA methylation (cg17401724) on gene expression (*ERBB2*), using the genotype at rs66826786 as the instrumental variable.

## Results

### Genome-wide *cis*-eQTLs and *cis*-meQTLs mapping in cultured airway epithelial cells

To identify genetic variation influencing gene expression under different conditions, we performed eQTL mapping in cultured AECs exposed to RV, and its corresponding vehicle control from 104 individuals (43 with doctor diagnosed asthma; 61 without a doctor’s diagnosis of asthma; Fig. S1). Analyses were performed separately for each treatment condition, testing for associations with 6,665,552 imputed SNPs (MAF>0.05) and 11,231 autosomal genes (see Methods; Additional file 1 and 2). The numbers of SNPs associated with gene expression for at least one gene (eQTLs) and genes with at least one eQTL (eGenes), in any treatment, are summarized in Fig. S5.

In parallel, we performed meQTL mapping in the same cells used for gene expression studies. We performed this analysis separately for each treatment condition, testing for associations with the same imputed SNP set that was used for eQTL mapping and interrogated 791,765 autosomal CpGs (Additional file 3 and 4). A summary of the number of SNPs associated with methylation levels at one or more CpG sites (meQTLs) and CpG sites with at least one meQTL (meCpGs), in any treatment, are shown in Fig. S5.

Each gene/CpG-variant pair was tested for a linear regression slope that significantly deviated from 0. Therefore, the estimated effects for the molecular QTLs reflects both the single-SNP effects of each molecular QTL as well as those that are in linkage disequilibrium (LD). Accordingly, these analyses do not differentiate between causal molecular QTLs from those in LD with the QTL. However, these variants are still informative in prioritizing genes and CpG sites that contribute to the etiology of asthma.

### Estimating shared and condition-specific molecular QTL effects

After identifying molecular QTLs in each treatment condition, we first explored the impact of RV exposure on eQTLs and meQTLs by comparing RV-treated to vehicle-treated results. For this analysis, we used an empirical Bayes method, multivariate adaptive shrinkage (*mash*; see the “Methods” section) [32]. Compared to direct comparisons between conditions, *mash* increases power, improves effect-size estimates, and provides better quantitative assessments of effect size heterogeneity of molecular QTLs, thereby allowing for greater confidence in effect sharing and estimates of condition-specificity. Additionally, as a confidence measurement of the direction of QTL effects, *mash* provides a ‘local false sign rate’ (lfsr) that is the probability that the estimated effect has the incorrect sign [39], rather than the expected proportion of Type I errors as would be assessed using FDR thresholds.

To identify condition-specific eQTLs, we analyzed the effect estimates of the most significant eQTL for each of 11,231 genes and assessed sharing of these signals among the RV and vehicle treated cells (see Methods). A pairwise comparison showed that 58.3% of eQTLs were shared between RV and vehicle treatments, representing 1,564 eGenes, defined here as genes with at least one eQTL at a lfsr < 0.05 (Fig. 1A; Additional file 5). We observed 660 and 458 condition-specific eGenes in the vehicle- and RV-treated cells, respectively. These potentially represent genetic variants that modify responses to viral exposure in AECs. Examples of treatment-specific eQTLs are shown in Fig. 1B. The effect estimates of the most significant meQTL for each of 751,914 CpG sites were used to identify condition-specific DNA methylation effects, as described above for eQTLs. A pair-wise analysis of meQTLs revealed that 89.9% of meQTLs were shared between vehicle and RV treatments, representing 48,189 meCpGs, defined here as CpGs with at least one meQTL at a lfsr<0.05 (Fig. 1C; Additional file 6), revealing a much greater proportion shared meQTLs than those observed for eQTLs. Examples of the 5,416 treatment-specific meQTLs are shown in Fig. 1D.

**Fig. 1.**
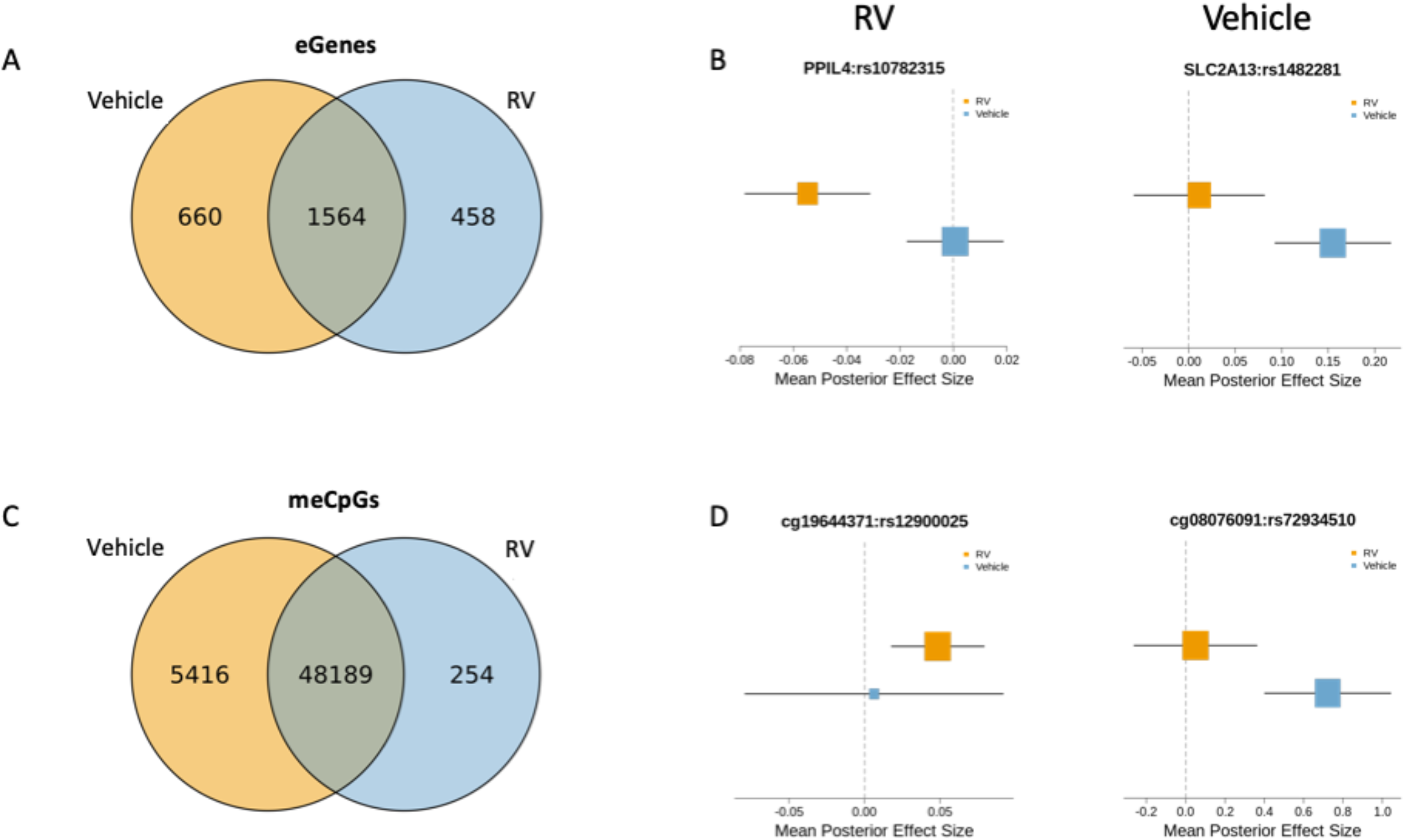
Summary of molecular effects sharing across treatment conditions (lfsr<0.05). Venn diagrams of eGenes **(A)** and meCpGs **(B)** shared between vehicle- and RV-treated airway epithelial cells. Forest plots showing examples of RV− (left) and Vehicle-specific (right) eQTLs **(C)** and meQTLs **(D)**.

In total, we identified 660 and 458 eGenes (lfsr<0.05) that were specific to vehicle or RV treatment, respectively, and 5,162 and 254 meCpGs that were specific to vehicle or RV culture treatment, respectively, with greater confidence than by pairwise comparisons using FDR thresholds [32].

### Molecular QTLs in the airway epithelium are enriched for asthma GWAS SNPs

Although the majority of variation in the human genome is non-functional [40], GWAS loci tend to be most enriched for functional annotations in disease-relevant cells [4, 41, 42]. To assess whether the 755,441 molecular QTLs identified in our study (i.e., the union of eQTLs and meQTLs from each treatment condition at FDR<0.10) are enriched for GWAS variants and whether these enrichments show tissue specificity, we first extracted summary statistics from a publicly available GWAS data for childhood onset and adult onset asthma [1] and for four diseases without known allergic or autoimmune etiologies (Alzheimer’s disease [34], atrial fibrillation [35], height [36], and neuroticism [37]). There were statistically significant enrichments (OR>1 and BH-adjusted P-value<0.05; see Methods) for the childhood and adult onset asthma GWAS SNPs among the molecular QTLs at each of four GWAS thresholds (Table 1), consistent with the strong epithelial cell involvement in asthma in general and with childhood onset asthma in particular. In contrast, there were no significant enrichments for SNPs from four of the other GWASs among the epithelial cell molecular QTLs. These results highlight the specific enrichment of asthma GWAS SNPs among airway epithelial molecular QTLs compared to SNPs from GWASs of diseases without known epithelial cells involvement.

**Table 1.**
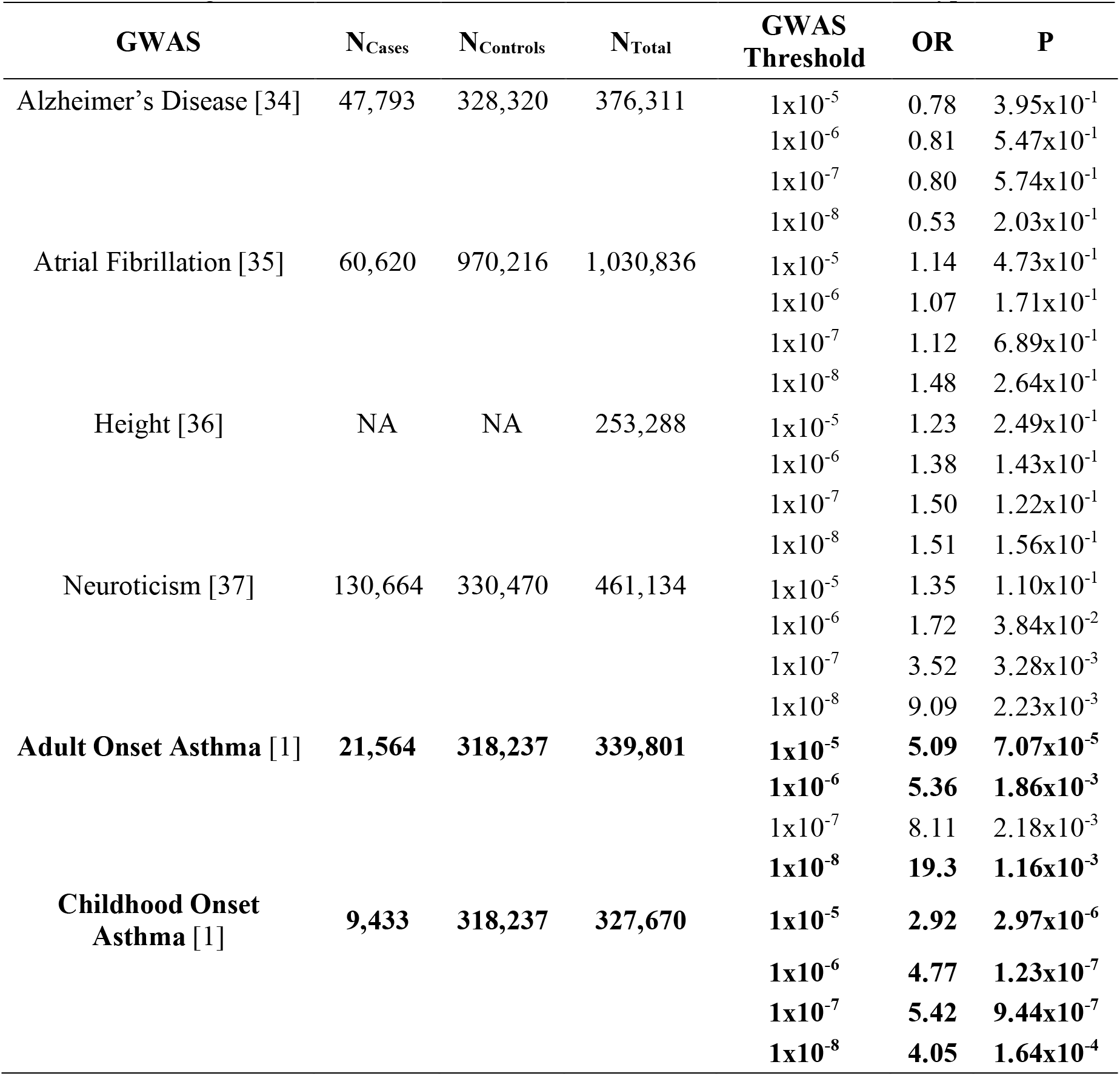
Enrichment estimates of airway epithelial cell molecular QTLs for GWAS SNPs. *P*-values that are significant after BH correction (see Methods) are shown in bolded type.

| GWAS | N<sub>Cases</sub> | N<sub>Controls</sub> | N<sub>Total</sub> | GWAS<br>Threshold | OR | P |
| --- | --- | --- | --- | --- | --- | --- |
| Alzheimer's Disease [34] | 47,793 | 328,320 | 376,311 | 1x10<sup>-5</sup> | 0.78 | 3.95x10<sup>-1</sup> |
|  |  |  |  | 1x10<sup>-6</sup> | 0.81 | 5.47x10<sup>-1</sup> |
|  |  |  |  | 1x10<sup>-7</sup> | 0.80 | 5.74x10<sup>-1</sup> |
|  |  |  |  | 1x10<sup>-8</sup> | 0.53 | 2.03x10<sup>-1</sup> |
| Atrial Fibrillation [35] | 60,620 | 970,216 | 1,030,836 | 1x10<sup>-5</sup> | 1.14 | 4.73x10<sup>-1</sup> |
|  |  |  |  | 1x10<sup>-6</sup> | 1.07 | 1.71x10<sup>-1</sup> |
|  |  |  |  | 1x10<sup>-7</sup> | 1.12 | 6.89x10<sup>-1</sup> |
|  |  |  |  | 1x10<sup>-8</sup> | 1.48 | 2.64x10<sup>-1</sup> |
| Height [36] | NA | NA | 253,288 | 1x10<sup>-5</sup> | 1.23 | 2.49x10<sup>-1</sup> |
|  |  |  |  | 1x10<sup>-6</sup> | 1.38 | 1.43x10<sup>-1</sup> |
|  |  |  |  | 1x10<sup>-7</sup> | 1.50 | 1.22x10<sup>-1</sup> |
|  |  |  |  | 1x10<sup>-8</sup> | 1.51 | 1.56x10<sup>-1</sup> |
| Neuroticism [37] | 130,664 | 330,470 | 461,134 | 1x10<sup>-5</sup> | 1.35 | 1.10x10<sup>-1</sup> |
|  |  |  |  | 1x10<sup>-6</sup> | 1.72 | 3.84x10<sup>-2</sup> |
|  |  |  |  | 1x10<sup>-7</sup> | 3.52 | 3.28x10<sup>-3</sup> |
|  |  |  |  | 1x10<sup>-8</sup> | 9.09 | 2.23x10<sup>-3</sup> |
| Adult Onset Asthma [1] | **21,564** | **318,237** | **339,801** | **1x10<sup>-5</sup>** | **5.09** | **7.07x10<sup>-5</sup>** |
|  |  |  |  | **1x10<sup>-6</sup>** | **5.36** | **1.86x10<sup>-3</sup>** |
|  |  |  |  | 1x10<sup>-7</sup> | 8.11 | 2.18x10<sup>-3</sup> |
|  |  |  |  | **1x10<sup>-8</sup>** | **19.3** | **1.16x10<sup>-3</sup>** |
| Childhood Onset<br>Asthma [1] | **9,433** | **318,237** | **327,670** | **1x10<sup>-5</sup>** | **2.92** | **2.97x10<sup>-6</sup>** |
|  |  |  |  | **1x10<sup>-6</sup>** | **4.77** | **1.23x10<sup>-7</sup>** |
|  |  |  |  | **1x10<sup>-7</sup>** | **5.42** | **9.44x10<sup>-7</sup>** |
|  |  |  |  | **1x10<sup>-8</sup>** | **4.05** | **1.64x10<sup>-4</sup>** |

To further assess the specificity of airway epithelial molecular QTLs to asthma, we compared GWAS SNP enrichments among the eQTLs in our study to those from tissues that are not known to be involved in asthma. To this end, we tested for enrichment of asthma GWAS SNPs among eQTLs (FDR<0.05) in five different tissues from the GTEx database (adrenal, frontal cortex, hypothalamus, ovary, testis) [43], and compared them to enrichments among the eQTLs from our study. We observed a significant enrichment (OR>1 and BH-adjusted P<0.05) of childhood onset asthma GWAS SNPs among the epithelial cell eQTLs at all GWAS P-value thresholds ≤1×10^−7^ (Table 2), while enrichments for adult onset asthma GWAS SNPs among the epithelial cell eQTLs were not observed at any GWAS threshold (Table S1). Except for the hypothalamus, which showed some enrichment at P < 10^−5^), no other enrichments of asthma GWAS SNPs were observed among eQTLs in other tissues, further supporting the specificity of our model and previous studies suggesting that epithelial barrier defects underlie risk for childhood onset, but not adult onset, asthma [1, 17, 18].

**Table 2.**
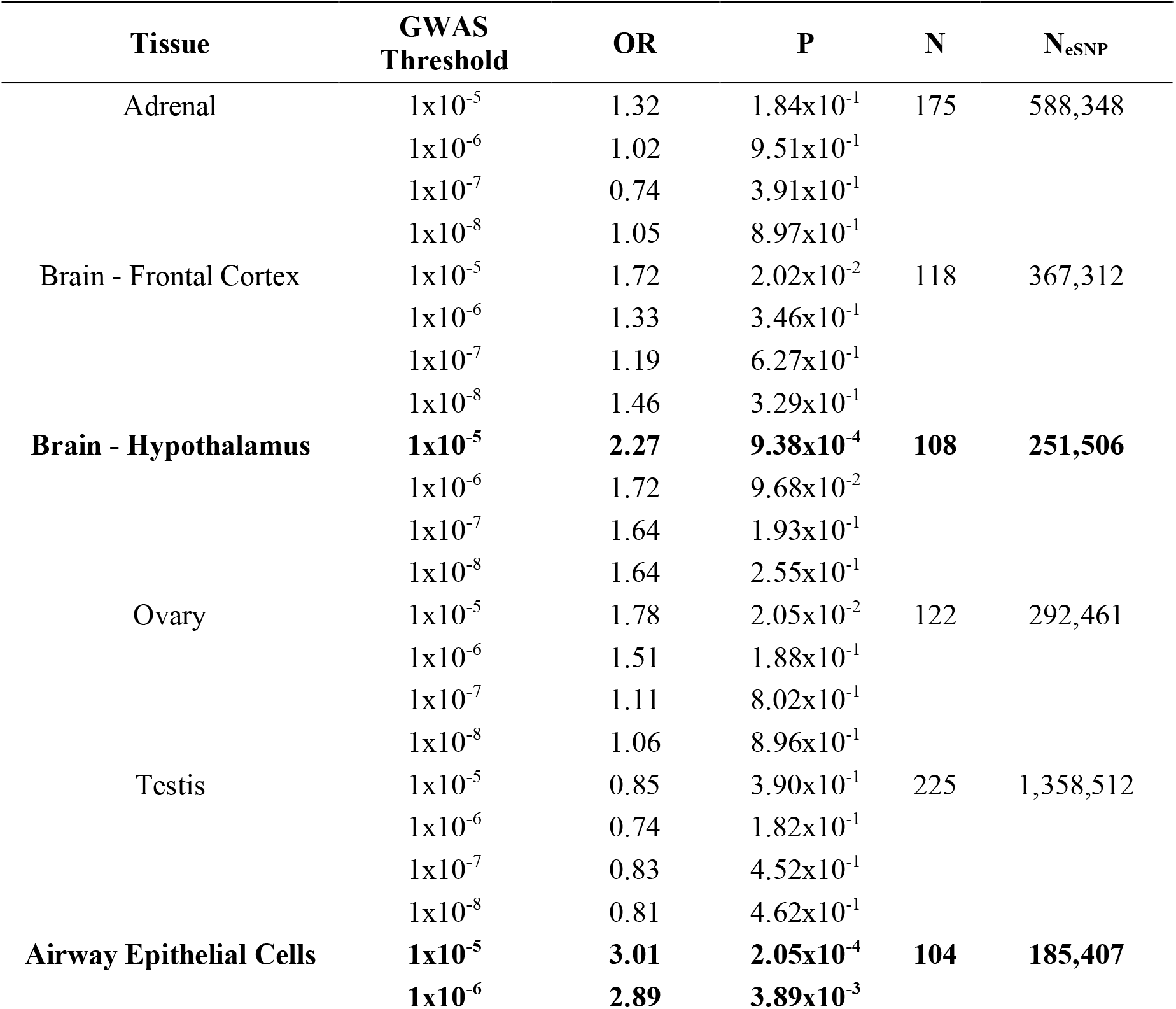
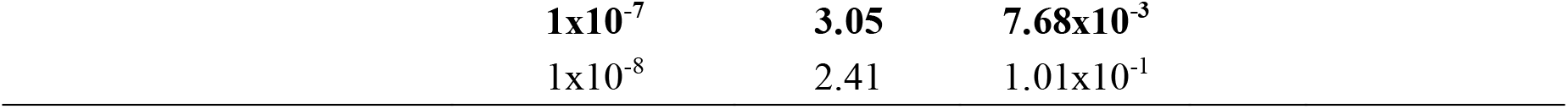
Enrichment estimates of eQTLs for childhood onset asthma GWAS SNPs from six tissues. Significant *P*-values after BH correction are shown in bolded type. Results for adult onset asthma is shown in Table S1.

### Molecular QTL co-localizations with adult onset and childhood onset asthma loci

Integrating molecular QTLs with GWAS data is a powerful way to identify functional variants that may ultimately influence disease risk [44, 45] and to assign function to known disease-associated variants. Co-localization approaches directly test whether the same genetic variant is underlying associations between two or more traits (e.g., gene expression and asthma), providing clues to causal disease pathways. We hypothesized that integrating molecular QTLs from RV- and vehicle-exposed epithelial cells with results of GWASs for adult onset and childhood onset asthma would reveal genetic and epigenetic mechanisms that modulate risk for childhood and/or adult onset asthma.

To test this hypothesis, we extracted summary statistics from large GWASs of adult onset asthma and childhood onset asthma [1], and tested each for co-localization with genetic variants associated with gene expression, DNA methylation, and asthma, using *moloc*, a Bayesian statistical approach that allows integration and co-localization of more than two molecular traits [13]. We performed four separate co-localization tests for each treatment conditions with each of the GWASs. Each analysis provided three possible configurations in which a variant is co-localized between the GWAS and QTLs: eQTL-GWAS pairs, meQTL-GWAS pairs, eQTL-meQTL-GWAS triplets. Estimates of a posterior probability of association (PPA) is provided, reflecting the evidence for a colocalized SNP being causal for the associations in the GWAS and for the corresponding eQTL and/or meQTL.

Using this approach, we found evidence for a total of 19 unique multiple trait co-localizations (Table 3). A single meQTL-GWAS pair was co-localized in both the adult onset and childhood onset asthma GWASs. An additional 18 co-localizations were detected only in the childhood onset asthma GWAS, including a single eQTL-meQTL-GWAS triplet associated with the *ERBB2* gene, three eQTL-GWAS pairs associated with three genes (*FLG, FLG-AS1, ORMDL3*), and 15 meQTL-GWAS pairs associated with 11 CpG sites (Table 3; Table S2). No co-localizations were specific to the adult onset asthma GWAS. Among the co-localized eGenes, based on previous studies, *FLG* was predicted to have decreased expression of in childhood onset asthma [46], *ERBB2* was predicted to have decreased expression of in severe asthma [47], and *ORMDL3* was predicted to have increased expression (Table S3) [48]. The larger number of co-localizations for childhood onset asthma relative to adult onset asthma is consistent both with the previous observation that genes at the childhood onset asthma loci were most highly expressed in skin, an epithelial cell type [1] and with the enrichment of childhood onset asthma GWAS SNPs among epithelial cell eQTLs described above.

**Table 3.** Number of QTL-GWAS pairs or triplets with evidence of co-localization (PPA≥0.70). The one meQTL-GWAS pair in the adult onset asthma GWAS is also among the 15 meQTL-GWAS pairs in the childhood onset GWAS.

| GWAS | eQTL-meQTL-GWAS | eQTL-GWAS | meQTL-GWAS |
| --- | --- | --- | --- |
| Adult onset asthma | 0 | 0 | 1 |
| Childhood onset asthma | 1 | 3 | 15 |
| PPA $\geq 0.70$ | | | |

The significance threshold (p<5×10^−8^) required to control the false discovery rate in GWASs likely excludes many true associations that do not reach this stringent cutoff. We and others have suggested that these SNPs, i.e., the mid hanging fruit [49], may be environment-or context-specific associations that are missed in GWASs that typically do not control for either [50, 51]. Notably, 10 of the 19 SNPs associated with co-localizations in the childhood onset asthma GWAS did not reach genome-wide significance in the GWAS. These results therefore provided functional inferences both for variants that were significant in a GWAS at known asthma loci and for variants that did not meet strict criteria for significance in the GWAS, thereby facilitating prioritization of variants among the mid-hanging fruit [49]. Two examples of co-localizations with prominent asthma-associated loci are described in the following sections.

### meCpGs at *TSLP* co-localize with an asthma risk variant

To more deeply characterize the co-localizations, we first focused on the only meQTL-GWAS pair in both the adult onset and childhood onset asthma GWASs. This pair included an intergenic SNP (rs1837253) located 5.7 kb upstream from the transcriptional start site (TSS) of the *TSLP* gene on chromosome 5q22, encoding an epithelial cell cytokine that plays a key role in the inflammatory response in asthma and other allergic diseases [52]. rs1837253 co-localized with a single meQTL (cg15557878) in both the adult onset (p_GWAS_ = 2.77×10^−13^) and childhood onset (p_GWAS_ = 2.33×10^−27^) asthma GWASs. The meCpG is located in the first (untranslated) exon (5’ UTR) of the *TSLP* gene (Fig. 2), a region characterized as a promoter in normal human epidermal keratinocyte cells (NHEK; ROADMAP). In fact, rs1837253 was the sentinel SNP at this locus in GWASs of asthma (e.g. [1, 53]) and of moderate-to-severe asthma [54]. In our study, the rs1837253-C asthma risk allele was associated with hypermethylation in primary cultured AECs at cg15557878 (Fig. 2), but was not associated with the expression of *TSLP* in either treatment condition (not shown).

**Fig. 2.**
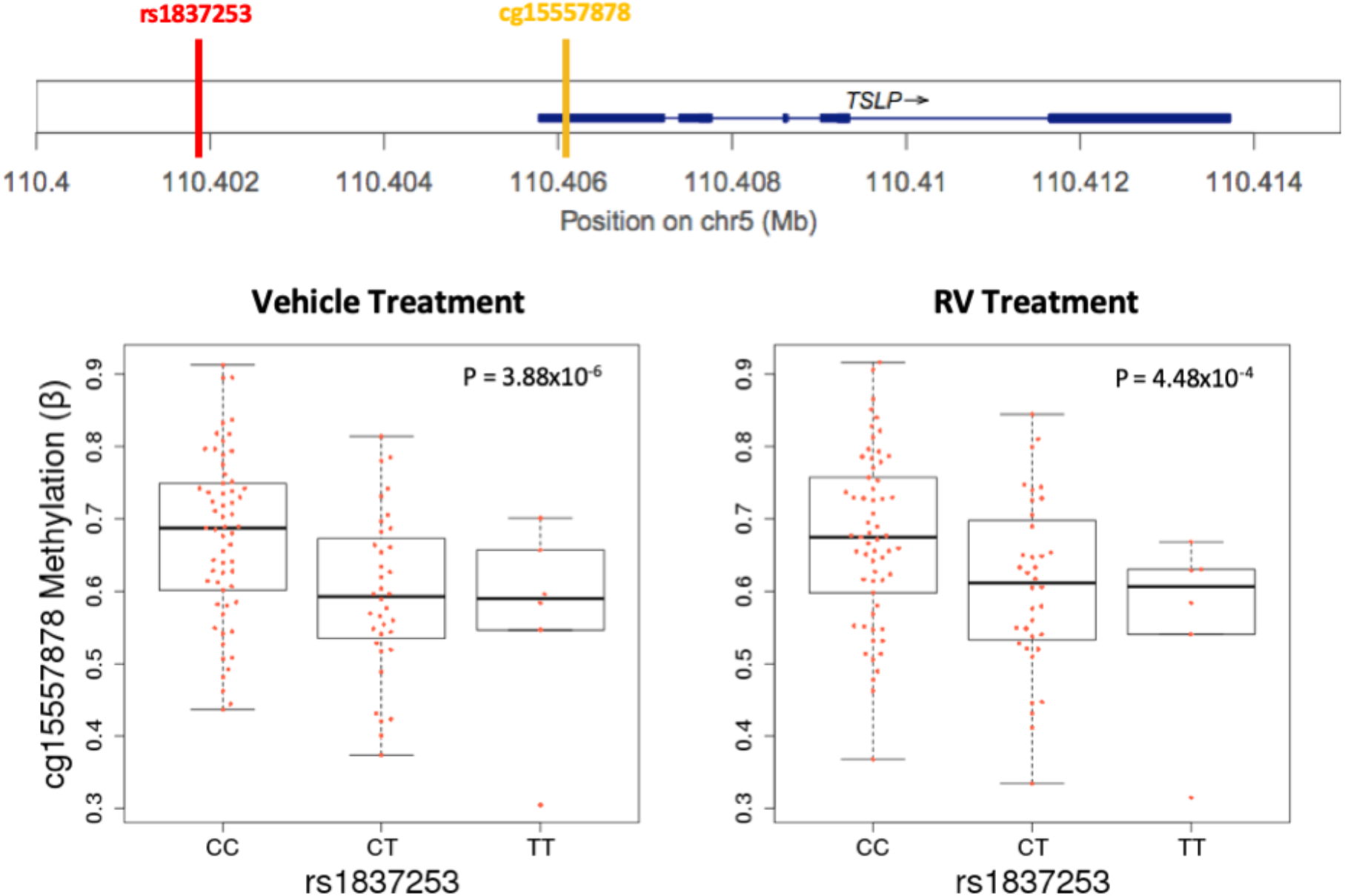
Co-localization of rs1837253 with DNA methylation levels for cg15557878 at *TSLP*. rs1837253 (red vertical bar, upper panel)) is associated with DNA methylation levels at cg15557878 (orange vertical bar, upper panel). Box plots show DNA methylation levels (y-axes) for each meCpGs by rs1837253 genotype (x-axes) in each treatment condition (lower panel).

Previous studies have shown *TSLP* to be a methylation-sensitive gene and that hypomethylation at its promoter is associated with atopic dermatitis (AD) and prenatal tobacco smoke exposure [55, 56]. Another study showed that the rs1837253-CC genotype was associated with increased excretion of TSLP in cultured AECs after exposure to polyI:C (a dsRNA surrogate of viral stimulation) [57]. Neither finding could be addressed in our study. Moreover, we were unable to identify any SNPs in LD with rs1837253 (± 50 kb) in either European or African American (*r*^2^ < 0.12) 1000Genomes reference panels, implying that this SNP may indeed be the causal SNP at this locus. Our results further suggest that DNA methylation levels in AECs may underlie this effect.

### Multi-trait co-localizations of molecular QTLs and asthma risk at the 17q12-21 asthma locus

To further explore the possibility that some mechanisms of asthma risk are exposure-specific, we focused on the co-localizations of eQTLs and meQTLs with asthma-associated SNPs at the 17q12-21 (17q) locus, the most replicated locus for childhood onset asthma (reviewed in [2]). This locus is characterized by high LD across a core region of 150 kb, encoding at least 4 genes (including *ORMDL3* and *GSDMB*). SNPs extending both proximal (including *PGAP3* and *ERBB2*) and distal (including *GSDMA*) to the core region show less LD with those in the core region and have been implicated as potentially independent asthma risk loci. Previous studies have shown that SNPs at this extended locus are eQTLs for at least four genes (*ORMDL3*, *GSDMB*, *GSDMA*, *PGAP3*) in blood and/or lung cells [2] and that genetic variants at this childhood onset asthma locus are also strongly associated with early life wheezing illness [58, 59], particularly RV-associated wheezing illness [60].

We identified six co-localizations at the extended 17q locus of molecular QTLs that were specific to childhood onset asthma GWAS SNPs. Among these co-localizations, one eQTL-GWAS pair with rs12603332 and expression of *ORMDL3* was only in vehicle-treated cells (PPA≥0.70; Fig. 3A-B). The co-localized SNP (rs12603332) is in LD (*r*^2^ > 0.74 in 1000 Genomes European reference panel) with other previously reported asthma-associated GWAS SNPs in this region, including some that were reported as eQTLs for *ORMDL3* and *GSDMB*, primarily in blood immune cells. However, in contrast to studies in *ex vivo* upper AECs [61], none of the SNPs were eQTLs for *GSDMB* in our *in vitro* culture model. That the co-localization with rs12603332 and *ORMDL3* expression was only significant in vehicle treated cells reflects the blunting of the eQTL effects (Fig. 3B), and possibly the overall decreased expression of *ORMDL3* (Fig. 3C), in RV-treated cells.

**Fig. 3.**
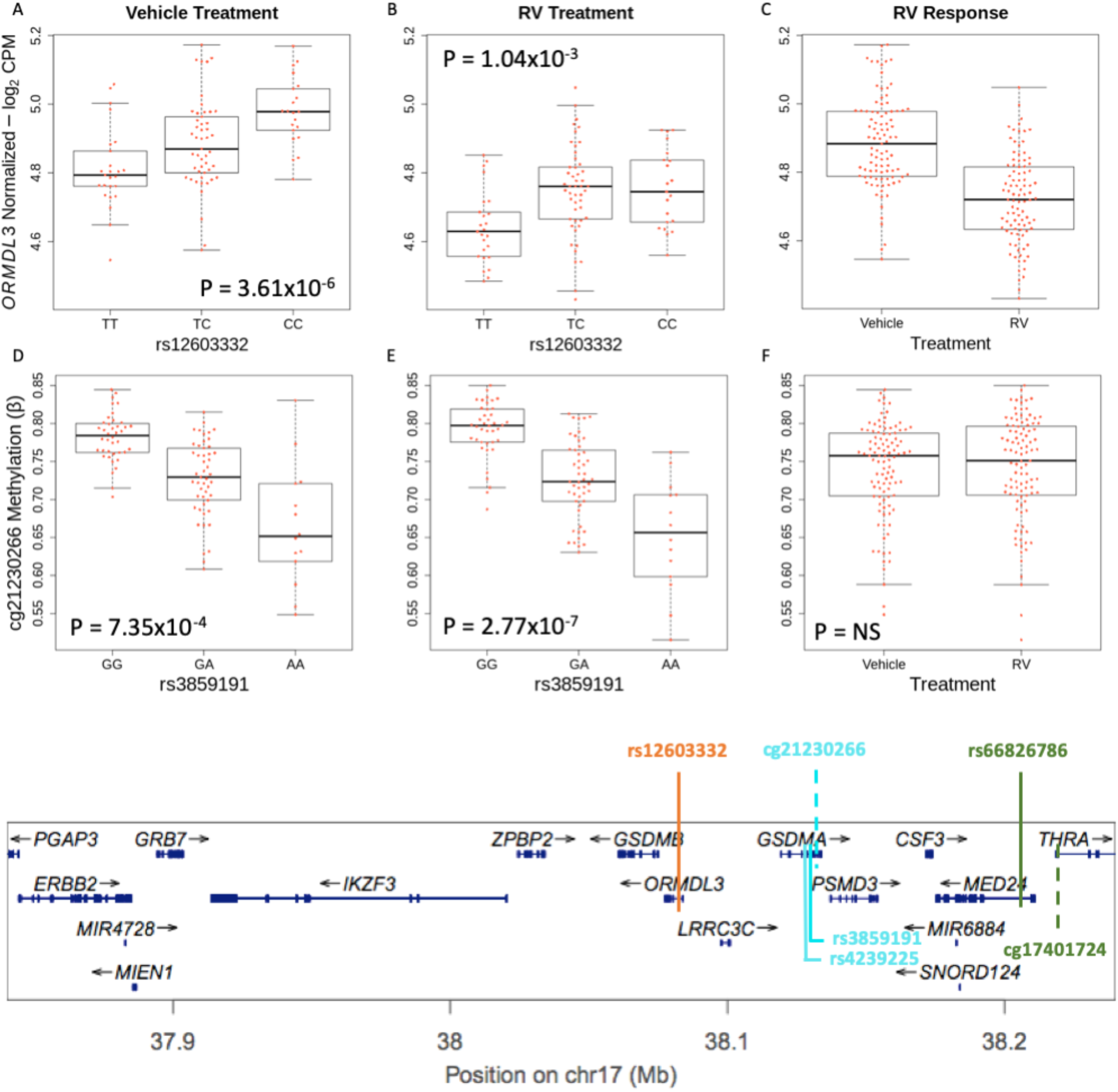
Co-localization pairs at the 17q asthma susceptibility locus. Upper panel: Box plots for the co-localized *ORMDL3* eQTL in cultured airway epithelial cells treated with vehicle **(A)** and RV **(B)**. The effect size of the correlation between *ORMDL3* and rs12603332 decreases after treatment with RV (FDR<0.10); and *ORMDL3* gene expression decreases after treatment with RV **(C)**. Box plots of the cg21230266 meQTL treated with vehicle (**D**) and RV (**E**), and methylation levels at cg21230266 (**F**). The meQTLs and overall methylation levels are similar in vehicle and RV treatments. Lower panel: The extended 17q12-21 locus. Co-localizations are shown by the vertical colored lines; rs4239225, a SNP in LD with rs3859191 (solid turquoise lines), is also significantly correlated with DNA methylation levels at cg21230266 in vehicle RV (not shown; see Table S2). See Fig 4B for box plots of the cg17401724:rs66826786 meQTL. Solid lines indicate the position of the colocalized SNP. Dashed lines indicate the location of meCpG pairs. Traits of the same co-localization are shown in the same color. A single eQTL-GWAS pair for *ORMDL3* is shown in orange; meQTL-GWAS pairs are shown in turquoise and green.

We also detected three meQTL-GWAS pairs among the six co-localizations at the 17q locus that were associated with two meCpGs (cg21230266, cg17401724) and three SNPs at the distal end of (rs4239225, rs3859191) and beyond (rs66826786) the extended locus near *GSDMA*, where there is some reduction of LD with SNPs in the core region (Fig. 3D-F). One of these CpGs was located in an intron (cg21230266) of *GSDMA* in regions characterized by ROADMAP as enhancers in NHEK cells. SNPs in modest to perfect LD (r^2^_range_=0.46 − 1.00; 1000 Genomes European panel) with these co-localizations (rs4239225, rs3859191) were described in previous studies as an independent GWAS signal for asthma (rs3894194) or an eQTL for *GSDMA* (rs3859192) [6, 62, 63]. These three meQTL-GWAS co-localizations were detected only in the RV-treated cells, although the meQTL signal for each of the three co-localizations was also detected in the vehicle treatment, likely due to decreased power to co-localize these meQTLs from the vehicle-treated cells. Additionally, there were no statistically significant differences in DNA methylation levels observed between the vehicle and RV treatments (Fig. 3F).

The one eQTL-meQTL-GWAS triplet detected in our study at the 17q locus (Fig. 4A, upper panel. The co-localization included an eQTL for *ERBB2*, at the proximal end of the locus and more than 361 kb from the co-localized asthma risk variant in an intron of *MED24* (rs66826786) and the co-localized meCpG (cg17401724) at the distal end of the locus (Fig. 3A, middle panel),. *MED24* is beyond the extended 17q12-21 locus as previously defined [2] in a region characterized by ROADMAP as both an enhancer and TSS in NHEKs. The eQTL for *ERBB2* is observed only after exposure to RV (Fig. 3A middle and lower panels), though the meQTL associated with this triplet was present in both vehicle and RV treatment conditions (Fig. 3B upper and lower panels, respectively). The asthma risk allele, rs66826786-T, was associated with decreased DNA methylation of cg17401724 in both conditions but with decreased *ERBB2* expression only in RV-treated cells. Overall, *ERBB2* expression decreased in response to RV exposure in AECs (Fig. 3D). The 361 kb distance between the promoter of *ERBB2* and its eSNP (rs66826786) suggests long-range interaction between *ERBB2* and the region harboring cg17401724 and rs66826786. The fact that the eQTL is observed only after RV infection, further suggests either that infection with RV triggers this long-range interaction in AECs via chromatin looping between these loci, or that RV infection results in the recruitment of negative transcription factors in this region that is already epigenetically poised. In fact, the observation that the meQTL for cg17401724 is observed in both conditions indeed suggests an epigenetically poised chromatin state at the distal end that directly affects transcription of *ERBB2* at the proximal end of the locus after exposure to RV, and possibly to other viruses.

**Fig. 4.**
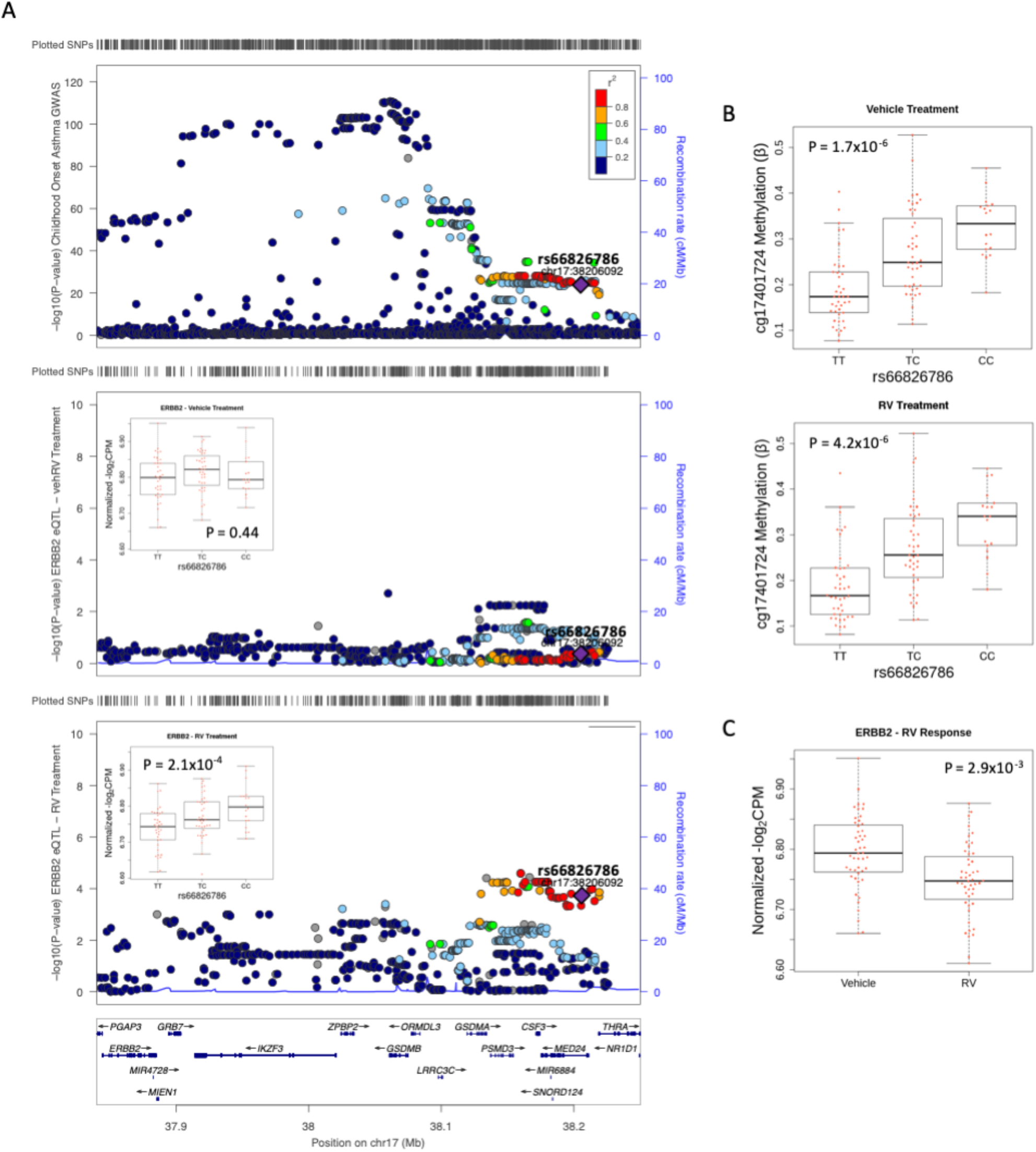
Co-localization of rs66826786 with *ERBB2* expression and DNA methylation levels for cg17401724. **(A)** LocusZoom plots of childhood onset GWAS results at the 17q locus showing the *ERBB2* gene at the proximal (left) end of the locus and the co-localized eQTL (rs668826786) at the distal (right) end of the locus (modified from Pividori ?). The SNP (rs66826786), which colocalized with associations for childhood onset asthma, *ERBB2* expression, and DNA methylation at cg17401724, is shown as a purple diamond in each of three LocusZoom plot. Upper panel: childhood onset asthma GWAS (modified from Pividori et al. 2019). Middle panel: *ERBB2* eQTLs for vehicle-treated cultured airway epithelial cells. Lower panel: *ERBB2* eQTLs for RV-treated airway epithelial cells. Boxplots for *ERBB2* gene expression by rs66826786 genotype is shown within the middle and lower LocusZoom plots. **(B)** Boxplots for cg17401724 meQTLs in vehicle-treated (upper panel) and RV-treated (lower panel) cultured airway epithelial cells. **(C)** *ERBB2* gene expression in vehicle-treated and RV-treated cells.

### Mendelian randomization of multi-trait co-localized triplets

Co-localization analyses reveal genetic variants that are associated with both asthma and molecular traits (gene expression and/or DNA methylation) but the question of causality between the molecular traits remains unanswered. To infer causal relationships between DNA methylation and gene expression on asthma risk, we performed Mendelian randomization (MR), a method in which genetic variation associated with modifiable exposure patterns (i.e. DNA methylation) can be used as an instrumental variable to estimate the causal influence of an exposure on an outcome (i.e. DNA methylation on gene expression) [64]. Specifically, we applied a two-stage least squares regression (2SLS; see Methods) regression to estimate the effects of DNA methylation (exposure) on gene expression (outcome) in each treatment condition, and used the QTL SNP (rs66826786) in the co-localized triplets at 17q as the genetic instrument (see Methods). In this way, we are able to estimate whether the effect of the asthma risk variant on gene expression levels is mediated by DNA methylation.

MR suggested a causal relationship between methylation and gene expression in RV-treated cells for the co-localized triplet, indicating that the genotype effect at rs66826786 on expression of *ERBB2* is mediated by methylation at the meCpG (Table 4). The contribution of the meCpG on *ERBB2* gene expression was only detected in RV-treated cells (P-value<1×10^−10^) while no evidence was detected in vehicle-treated cells (P-value=0.81), further suggesting a gene regulatory mechanism that is triggered after exposure to RV. The MR result provides orthogonal evidence for the co-localization of this triplet and novel evidence for causal inference with respect to the co-localized traits (DNA methylation, gene expression). These data also reinforce arguments for epigenetic mechanisms modifying gene expression, and potentially disease risk, in response to environmental exposures [65, 66].

## Discussion

One of the major challenges of complex disease genetics is to uncover molecular mechanisms of pathogenesis and to understand how genetic and environmental factors interact to influence risks for disease. While GWASs have identified thousands of SNPs associated with disease phenotypes, interpretation and downstream follow-up studies of GWAS results have been limited. Cell models can advance our understanding of disease pathobiology through experimental testing of disease mechanisms in a controlled environment. In this multi-omics study, we leveraged an airway epithelial cell model of microbial response to identify potentially functional variants, some of which have context-specific effects on transcriptional and epigenetic responses, and molecular mechanisms of disease. We show that asthma GWAS SNPs were specifically enriched among molecular QTLs in airway epithelial cells compared to SNPs from other GWASs, and among AEC eQTLs compared to eQTLs from other tissues. Finally, SNPs that were molecular QTLs in our study co-localized with asthma GWAS SNPs, identifying 18 unique co-localizations that included both known asthma loci (e.g., 17q12-21 and *TSLP*) and loci that did not meet stringent criteria for genome-wide significance in the GWASs (Table S2).

The results of enrichment analyses further highlighted the important role of airway epithelium in asthma GWAS discoveries. The enrichment of childhood onset asthma GWAS SNPs among epithelial eQTLs is particularly noteworthy, as it not only supports the tissue specificity of our model but also identified genomic loci with molecular mechanisms that have not been described prior to our study. These results are also consistent with previous studies suggesting that functional variants from disease-relevant tissues are more enriched among GWAS loci for those diseases [4, 41, 42]. The more modest enrichment of adult onset asthma GWAS SNPs among epithelial eQTLs may be due to the overall smaller effect sizes of SNPs at adult onset asthma loci compared to childhood onset asthma loci, to the less important role of epithelial cells in the pathophysiology of adult onset asthma, or to the greater heterogeneity and lesser heritability of adult onset asthma [1]. Other differences between the adult onset and childhood onset asthma GWASs were observed. For example, only a single co-localization was detected with adult onset asthma GWAS SNPs, compared to 19 with childhood onset asthma GWAS SNPs. None of the co-localizations in the adult onset GWAS included an eQTL compared to four childhood onset co-localizations with eQTLs, and the one meQTL-GWAS pair in the adult onset asthma GWAS was also present in the childhood onset asthma. These differences were additionally surprising because although there were 2.5-times the number of loci associated with childhood onset asthma compared to adult onset asthma in the GWASs [1], there were nearly 20-times more co-localizations in the childhood onset compared to the adult onset GWAS (19 vs. 1, respectively). These observations likely reflect the more important role of gene regulation and dysregulation in airway epithelium in the etiology of childhood onset asthma compared to adult onset asthma [17, 18]. Focusing on other asthma relevant tissues (e.g., lung tissue) or cells (e.g., immune cells) might reveal additional novel molecular mechanisms and differences between childhood onset and adult onset asthma.

Our study provides mechanistic evidence for associations between GWAS SNPs and asthma at two important loci: the *TSLP* and 17q12-21 loci. Co-localizations of the asthma associated SNP rs1837253 with DNA methylation levels in the *TSLP* gene suggest an epigenetic mechanism of disease that contributes to both adult and childhood onset asthma, and is robust to RV versus vehicle treatment. Associations of this SNP with asthma have been highly replicated in GWASs, and TSLP is recognized as having an important role in asthma pathogenesis through its broad effects on innate and adaptive immune cells promoting Th2 inflammation [67]. Our data further show that the effect of rs1837253 genotype on risk for asthma may be mediated through DNA methylation levels at CpG sites in the untranslated first exon of the *TSLP* gene in AECs. Finally, the lack of LD with other SNPs in a 100 kb window suggests that rs1837253 may be the causal asthma SNP at this important locus.

Since its discovery over a decade ago, the 17q12-21 locus has been an important focus of asthma research. Several studies have revealed the complex nature of this locus including the differences in LD structure across populations, and contrasting gene expression patterns and eQTLs at this locus in asthma-relevant cell types (reviewed in [2, 61]). In our study, using an airway epithelium cell model of RV infection, additional dimensions of complexity at this locus were revealed. For example, genes in the core region have been considered the most likely candidate genes mediating effects of genetic variation on risk of childhood onset asthma [61]. However, our study further shows that genes at both the proximal and distal ends of this locus, *ERBB2* and *GSDMA*, respectively, may contribute to asthma risk in the presence of RV infection. Mendelian randomization revealed a novel epigenetic mechanism through which a SNP at the distal boundary of the locus was associated with expression of *ERBB2* at the proximal boundary of the locus, only after exposure to RV. The eQTL effect on *ERBB2* expression in RV-treated cells was mediated through differential methylation of a CpG site at the distal locus, which was present in both treatment conditions. Previous studies have shown that variation at the 17q core locus confers risk to asthma only among children with wheezing illness in early life [68], particularly with RV-associated wheezing [59, 60]. Our study further connected RV infection and genotype at this locus to the *ERBB2* gene for the first time, as well as to an interaction between genetic and methylation variation at the distal end of the locus with the expression of *ERBB2* at the proximal end of the locus in RV infected epithelial cells. The SNP that is the eQTL for *ERBB2* in RV infected epithelial cells was associated with childhood onset asthma (p_GWAS_ = 6.43×10^−26^ [1]), directly connecting the eQTL for *ERBB2* in RV-treated cells to asthma risk. The asthma associated allele, rs66826786-T, was associated with decreased expression of *ERBB2* after RV infection in our study (Fig. 3A), consistent with results of a study of 155 asthma cases and controls reporting an inverse correlation between *ERBB2* expression in *ex vivo* lower AECs and asthma severity [47]. These combined data suggest that decreased expression of *ERBB2* associated with asthma severity may be modulated by RV, the most common trigger of asthma exacerbations, via epigenetic mechanisms involving DNA methylation and long-range chromatin looping between the proximal and distal ends of this important locus. In addition, meQTLs in *GSDMA*, at the proximal end of the locus, co-localized with GWAS SNPs in RV-treated cells only. Together, these findings further highlight the importance of RV exposure at this prominent asthma risk locus and provide mechanistic evidence for a genotype by exposure interaction, and raise the possibility that SNPs in the core region primarily confer risk for inception of early onset asthma whereas SNPs in the proximal and distal ends of the locus primarily modulate gene-environment interactions.

Many of the associations in GWASs that do not reach stringent criteria for genome-wide significance (p<5×10^−8^) may be true signals. Distinguishing true from false positive signals for variants among the mid-hanging fruit (e.g., p-values between 10^−5^ and > 10^−8^) can be challenging. In our study, over 57% of the co-localizations were with a GWAS SNP that did not meet genome-wide significance (childhood onset asthma GWAS p-value range 6.1×10^−7^ – 1.4×10^−5^; Table S2). One possibility for this is because the variants have exposure-specific, tissue-specific, or endotype-specific effects, which are heterogeneous among subjects included in GWASs. Therefore, annotating SNPs among the mid-hanging fruit for functionality provides more confidence to these findings, a more complete picture of the genetic architecture of asthma, and a model for prioritizing these loci for further studies.

Our study has several limitations. First, the sample sizes for the eQTL and meQTL studies were smaller than the most reliable sample size recommended by moloc (n_min_=300) [13]. In such cases, *moloc* can miss true co-localizations in QTL datasets. For example, an eQTL-GWAS pair with supporting evidence may, in reality, be an eQTL-meQTL-GWAS triplet. As a result, the eQTL-GWAS and meQTL-GWAS pairs that we identified could be eQTL-meQTL-GWAS triplets that we were not powered to detect, or we may have missed other co-localizations entirely. For example, although only a single meQTL co-localized with a GWAS SNP at the *TSLP* locus, the same SNP, rs1837253, was an meQTL for three additional CpGs (Fig. S6), representing additional potential contributors to asthma disease mechanisms. Nonetheless, the 19 unique co-localizations detected in our study are likely to be real, although future studies in larger samples will increase confidence in our findings. Second, we focused our studies on one cell type (upper airway sinonasal epithelium), two exposures (vehicle and RV), and one epigenetic mark (DNA methylation). It is possible that other asthma-relevant co-localizations are specific other tissues or cell types or to other exposures or culture conditions, and that additional epigenetic marks, such as those associated with chromatin accessibility, would be additionally informative. These extended studies will be necessary to validate the specificity and provide a more complete catalog of asthma-relevant co-localizations. Finally, characterizing chromatin conformational changes in AECs before and after exposure to RV will allow a direct assessment of the chromatin looping at the extended 17q12-21 locus that may occur in response to viral infection and potentially identify other context-specific interactions.

In summary, we identified *cis*-eQTLs and *cis*-meQTLs in an airway epithelial cell model of host cell response to RV and integrated those data with asthma GWASs to assign potential molecular mechanisms for variants associated with asthma in two large GWASs. By combining enrichment studies, co-localization analysis, and Mendelian randomization, we provide robust statistical evidence of epigenetic mechanisms in upper airway cells contributing to childhood onset asthma. We demonstrate that a multi-omics approach using a disease-relevant cell type and disease-relevant exposure allows prioritization of disease-associated variants and provides insight into potential epigenetic mechanisms of asthma pathogenesis.

## Acknowledgments

The authors acknowledge Christine Billstrand and Raluca Nicolae for sample processing and library preparation, and study subjects for their participation. This work was supported by NIH grants U19 AI106683 and R01 HL129735. M.M.S. was supported in part by T32 GM007197.

## Supporting information

**Fig. S1** Overview of the e/meQTL and co-localization studies in NECs treated with RV. **(A)** Step-wise experimental design to identify treatment-specific e/meQTLs in NECs from 104 individuals: 1. NECs collected from study participants were cultured and treated with RV and a vehicle for 48 hours. 2. Gene expression and DNA methylation measured in NECs from each treatment condition. 3. Genotype profiling to identify genetic variation influencing gene expression and DNA methylation to RV- and SA-treatment. 4. QC and analyses including e/meQTL mapping, multi-trait co-localization analysis, and Mendelian randomization. **(B)** Breakdown of the number of subjects for each experiment and molecular QTL mapping.

**Fig. S2 PCA and k-means clustering of genotypes. (A)** PCA plot of study participant’s genotypes (circles) projected on HapMap genotypes (squares). **(B)** Scree plot of k-means clustering of ancestral PCs in which the within groups sum of squares (y-axis) is plotted against the number of potential group clusters (x-axis); using the ‘elbow criterion’, it is determined that two clusters are best representative of how many clusters study samples can be grouped into. **(C)** PCA plot of study participants grouped into two cluster for genotype imputation, European (red), and African American (Blue), according to the k-means clustering criterion.

**Fig. S3** PCA of gene expression in RV- and vehRV-treated epithelial cells. **(A)** PCA plot of epithelial cell gene expression from 95 individuals treated with vehicle and RV before regressing out covariates. **(B)** PCA plot of gene expression in vehicle- and RV-treated cells after regressing out covariates. Tables showing p-values of correlation with PCs and covariates before **(C)** and after **(D)** regression. **(E)** Volcano plot showing differential gene expression in response to RV treatment.

**Fig. S4** PCA of DNA methylation in vehicle- and RV-treated in cultured airway epithelial cells. **(A)** PCA plot of cultured airway epithelial DNA methylation from 103 individuals treated with vehicle and RV before regressing out covariates. **(B)** PCA plot of DNA methylation in vehicle- and RV-treated cells after regressing out covariates. Tables showing p-values of correlation with PCs and covariates before **(C)** and after **(D)** regression. **(E)** Volcano plot showing differential DNA methylation in response to RV treatment.

**Fig. S5** Summary results for molecular QTL mappings. Venn diagrams of eQTLs **(A)** and meQTLs **(B)** in each condition (FDR<0.10). **(C)** Summary of eQTL and meQTL mapping results for each treatment condition. The number of SNPs associated with the gene expression of at least one gene or CpG and the number of genes or CpGs whose expression or DNA methylation levels was associated with at least one SNP.

**Fig. S6** meQTLs at rs1837253 located in the first untranslated exon of the *TSLP* gene. Box plots of three meQTLs that were identified in both the vehicle- (left) and RV-treated (right) AECs were correlated with the asthma risk variant (rs1837253) but did not show evidence of co-localization. These meQTLs may represent additional evidence of an epigenetic mechanism contributing to asthma.

**Table S1** Enrichment estimates of eQTLs for adult onset asthma GWAS SNPs from six tissues. P-values that are significant after BH correction are shown in bolded type.

**Table S2** *moloc* results indicating molecular QTL-GWAS pars and triplets

**Table S3** Adult onset and childhood onset asthma GWAS risk allele effects on gene expression and DNA methylation

