## Supplemental Figures for "Multi-omics co-localization with genome-wide association studies reveals a context-specific genetic mechanism at a childhood onset asthma risk locus"

S1 Figure

A

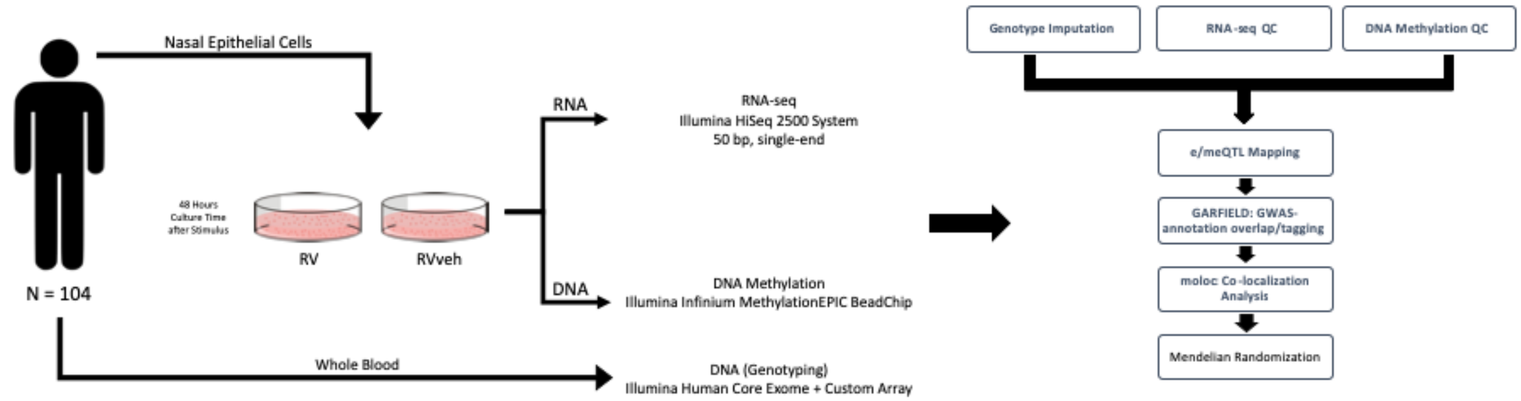


B

**Experimental sample composition**

| QTL Study | cis-eQTL | cis-meQTL |
|---|---|---|
| Sample Size (N) | 95 | 103 |
| Gender (% Female) | 45 (18 - 73) | 45 (18 - 73) |
| Mean Age (Range) | 41 | 33 |
| % Asthma (Ever) | 44 | 41 |

#### S2 Figure

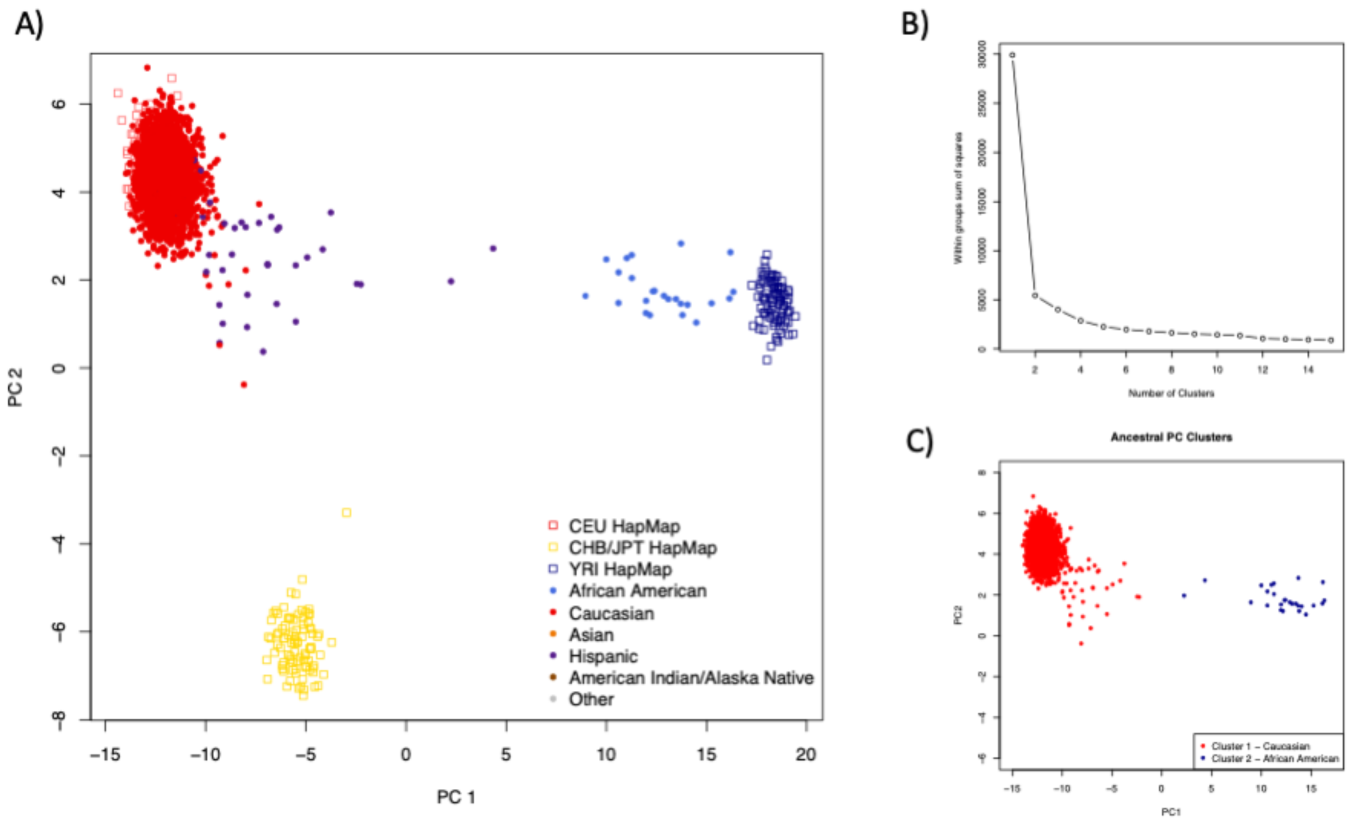
A)
B)
C)
PC 1
PC 2
CEU HapMap
CHB/JPT HapMap
YRI HapMap
African American
Caucasian
Asian
Hispanic
American Indian/Alaska Native
Other
Within groups sum of squares
Number of Clusters
Ancestral PC Clusters
Cluster 1 – Caucasian
Cluster 2 – African American
PC1
PC2

### S3 Figure

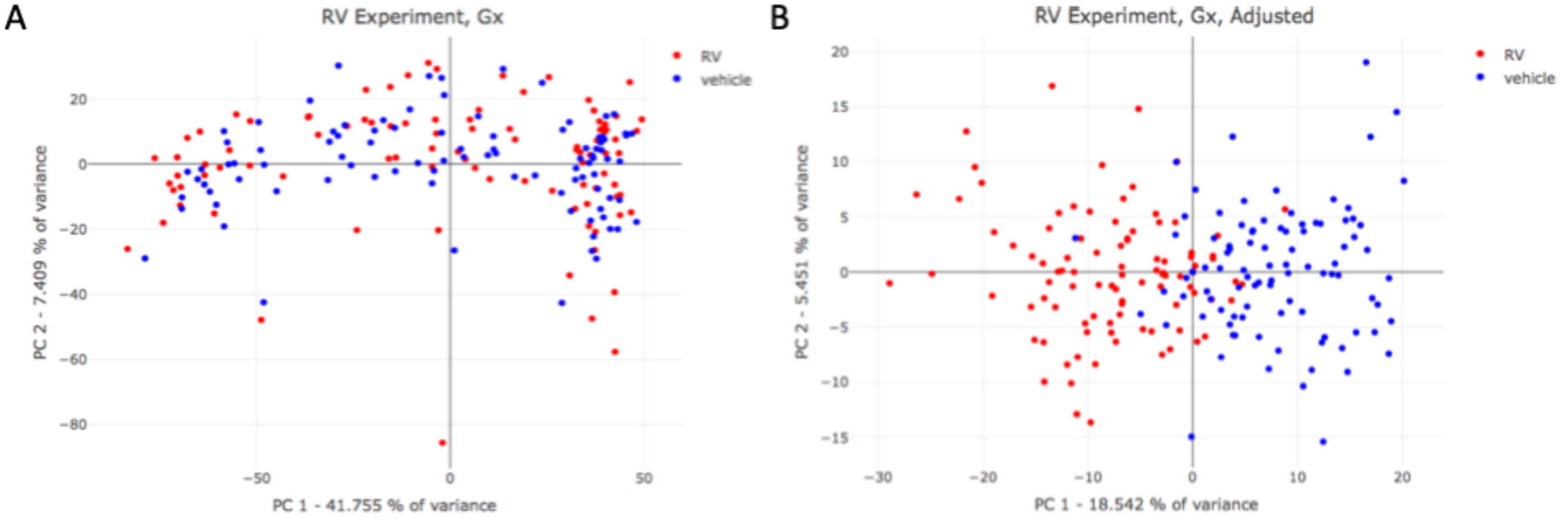


C

| PC (Variance Explained) | Treatment (RV) | Sex | Day in Liquid Nitrogen | Cell Cult. Batch | Days Exp. Cult. | RNA Conc. | RNA Frag. Length | Tech. | Seq. Pool | Ancestral PC 1 | Ancestral PC 2 | Ancestral PC 3 |
|---|---|---|---|---|---|---|---|---|---|---|---|---|
| PC1 (41.8%) | 9.51E-01 | 9.64E-01 | 6.28E-05 | 1.86E-15 | 5.59E-01 | 1.37E-38 | 1.41E-01 | 4.56E-01 | 3.20E-03 | 2.18E-02 | 7.75E-01 | 4.15E-01 |
| PC2 (7.41%) | 8.81E-01 | 7.09E-06 | 5.18E-01 | 1.34E-04 | 3.50E-01 | 4.72E-01 | 2.37E-02 | 6.55E-09 | 1.82E-01 | 7.36E-02 | 6.24E-01 | 2.11E-02 |
| PC3 (6.89%) | 6.52E-03 | 1.11E-01 | 8.42E-01 | 2.06E-01 | 5.35E-09 | 8.20E-04 | 1.74E-01 | 9.92E-01 | 3.04E-01 | 1.33E-01 | 3.21E-01 | 1.87E-01 |
| PC4 (5.74%) | 1.17E-09 | 2.35E-01 | 5.64E-01 | 2.82E-04 | 3.38E-01 | 9.93E-01 | 3.35E-02 | 1.46E-01 | 1.54E-01 | 3.99E-03 | 2.35E-01 | 6.52E-01 |
| PC5 (3.97%) | 4.48E-07 | 2.09E-02 | 2.65E-01 | 1.50E-03 | 2.35E-01 | 4.46E-01 | 6.98E-07 | 7.78E-01 | 3.52E-03 | 1.50E-01 | 2.25E-05 | 6.81E-01 |
| PC6 (3.55%) | 2.82E-01 | 2.59E-02 | 1.22E-01 | 9.92E-08 | 3.71E-02 | 4.66E-01 | 4.70E-01 | 2.36E-01 | 8.01E-02 | 5.03E-01 | 1.57E-01 | 2.33E-04 |
| PC7 (2.47%) | 5.33E-08 | 4.96E-02 | 2.19E-02 | 2.87E-04 | 6.31E-02 | 1.00E-01 | 7.68E-01 | 8.67E-01 | 7.63E-01 | 4.05E-04 | 7.62E-04 | 2.81E-01 |
| PC8 (2.06%) | 2.08E-01 | 4.23E-02 | 7.64E-01 | 5.93E-01 | 3.72E-04 | 7.03E-01 | 4.97E-01 | 2.81E-01 | 3.73E-02 | 1.02E-03 | 2.63E-02 | 6.19E-02 |
| PC9 (1.49%) | 7.23E-02 | 1.92E-01 | 1.05E-01 | 1.08E-05 | 9.69E-04 | 3.02E-01 | 7.34E-01 | 1.09E-01 | 2.69E-01 | 5.62E-04 | 2.65E-03 | 6.32E-03 |
| PC10 (1.07%) | 1.95E-02 | 7.59E-04 | 2.81E-01 | 1.19E-01 | 7.16E-01 | 2.27E-01 | 7.89E-01 | 6.26E-03 | 4.98E-04 | 2.55E-03 | 9.65E-01 | 9.52E-01 |

D

| PC (Variance Explained) | Treatment (RV) | Sex | Day in Liquid Nitrogen | Cell Cult. Batch | Days Exp. Cult. | RNA Conc. | RNA Frag. Length | Tech. | Seq. Pool | Ancestral PC 1 | Ancestral PC 2 | Ancestral PC 3 |
|---|---|---|---|---|---|---|---|---|---|---|---|---|
| PC1 (18.5%) | 1.91E-36 | 1 | 1 | 1 | 1 | 1 | 1 | 1 | 1 | 1 | 1 | 1 |
| PC2 (5.45%) | 8.38E-01 | 1 | 1 | 1 | 1 | 1 | 1 | 1 | 1 | 1 | 1 | 1 |
| PC3 (4.10%) | 2.66E-02 | 1 | 1 | 1 | 1 | 1 | 1 | 1 | 1 | 1 | 1 | 1 |
| PC4 (3.32%) | 5.27E-01 | 1 | 1 | 1 | 1 | 1 | 1 | 1 | 1 | 1 | 1 | 1 |
| PC5 (2.92%) | 7.74E-01 | 1 | 1 | 1 | 1 | 1 | 1 | 1 | 1 | 1 | 1 | 1 |

E

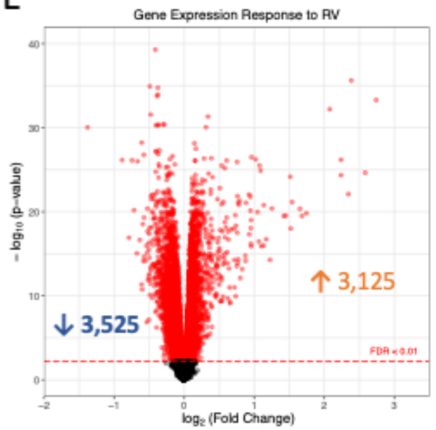

#### S4 Figure

A

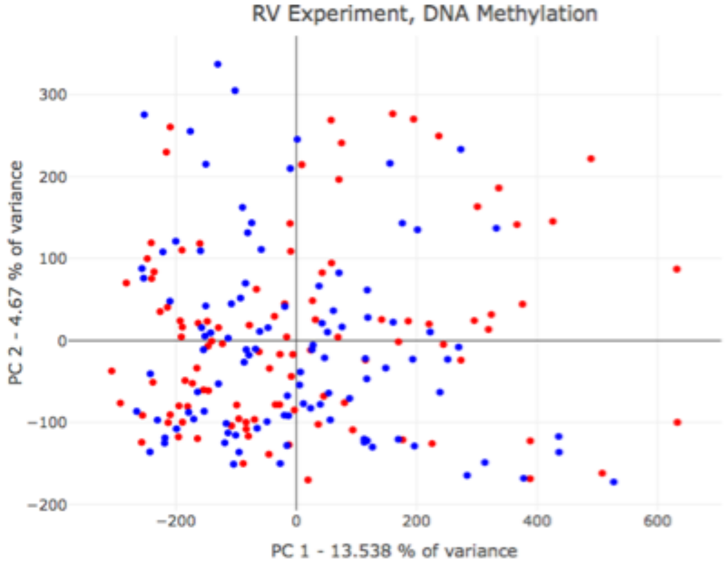


B

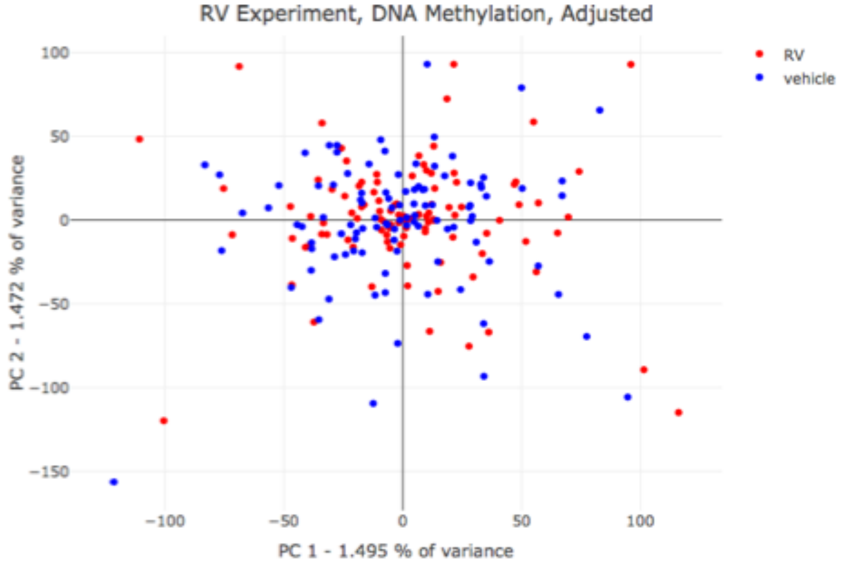


C

| PC (Variance Explained) | Treatment (RV) | Sex | Age | Smoking Status | Array | Cell Harvest Date |
|---|---|---|---|---|---|---|
| PC1 (13.5%) | 8.57E-01 | 3.80E-01 | 6.78E-01 | 2.95E-02 | 4.53E-01 | 3.68E-04 |
| PC2 (4.67%) | 4.28E-01 | 1.61E-01 | 8.65E-01 | 2.40E-01 | 6.04E-01 | 5.44E-06 |
| PC3 (3.83%) | 6.27E-02 | 7.26E-01 | 4.30E-02 | 5.05E-01 | 8.42E-05 | 2.08E-01 |
| PC4 (3.52%) | 5.19E-01 | 8.96E-02 | 7.90E-03 | 1.42E-01 | 6.54E-01 | 8.75E-06 |
| PC5(2.63%) | 7.55E-01 | 8.27E-01 | 1.09E-01 | 3.06E-01 | 2.66E-01 | 6.32E-03 |
| PC6 (2.01%) | 5.82E-01 | 8.96E-01 | 2.90E-14 | 4.27E-04 | 5.76E-01 | 8.99E-10 |
| PC7 (1.73%) | 5.28E-01 | 1.03E-04 | 7.74E-01 | 6.55E-01 | 2.79E-01 | 2.43E-03 |
| PC8 (1.47%) | 9.94E-01 | 6.98E-03 | 1.14E-01 | 7.56E-01 | 1.23E-01 | 2.30E-02 |
| PC9 (1.29%) | 7.23E-01 | 5.24E-01 | 5.70E-03 | 9.41E-01 | 2.94E-01 | 1.38E-04 |
| PC10 (1.18%) | 1.63E-02 | 4.93E-02 | 6.78E-01 | 2.95E-02 | 4.53E-01 | 3.68E-04 |

D

| PC (Variance Explained) | Treatment (RV) | Sex | Age | Smoking Status | Array | Cell Harvest Date |
|---|---|---|---|---|---|---|
| PC1 (1.56%) | 4.98E-01 | 1 | 1 | 1 | 1 | 1 |
| PC2 (1.50%) | 6.84E-01 | 1 | 1 | 1 | 1 | 1 |
| PC3 (1.49%) | 2.40E-01 | 1 | 1 | 1 | 1 | 1 |
| PC4 (1.45%) | 6.30E-01 | 1 | 1 | 1 | 1 | 1 |
| PC5 (1.45%) | 5.80E-01 | 1 | 1 | 1 | 1 | 1 |

E

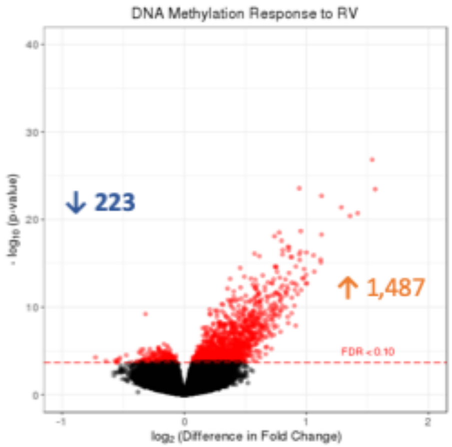

**S5 Figure**

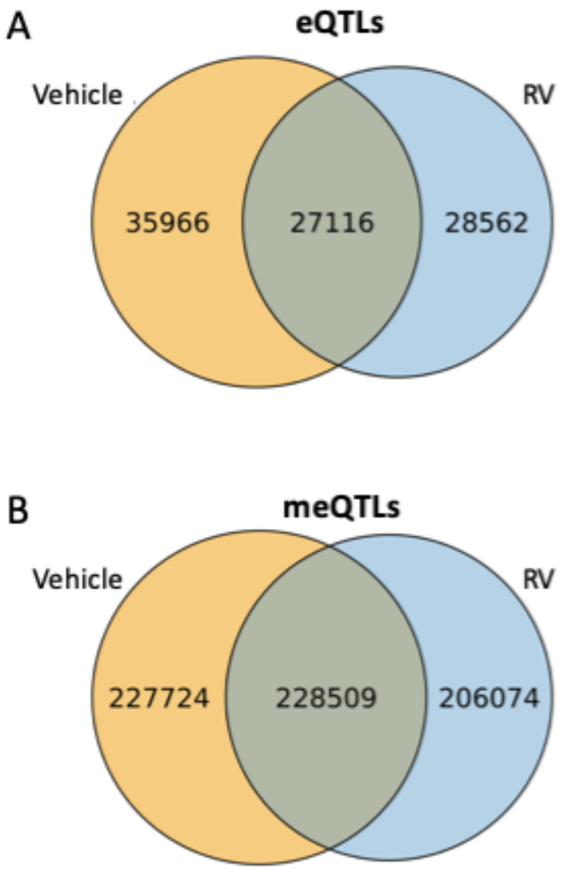


C

eQTL and meQTL mapping results (FDR<0.10)

| | Vehicle | RV |
|---|---|---|
| **eQTLs** | 63,082 | 55,678 |
| SNPs | 59,689 | 52,519 |
| Genes | 1,873 | 1,637 |
| **meQTLs** | 456,233 | 434,483 |
| SNPs | 320,174 | 306,850 |
| CpGs | 42,038 | 40,789 |

### S6 Figure

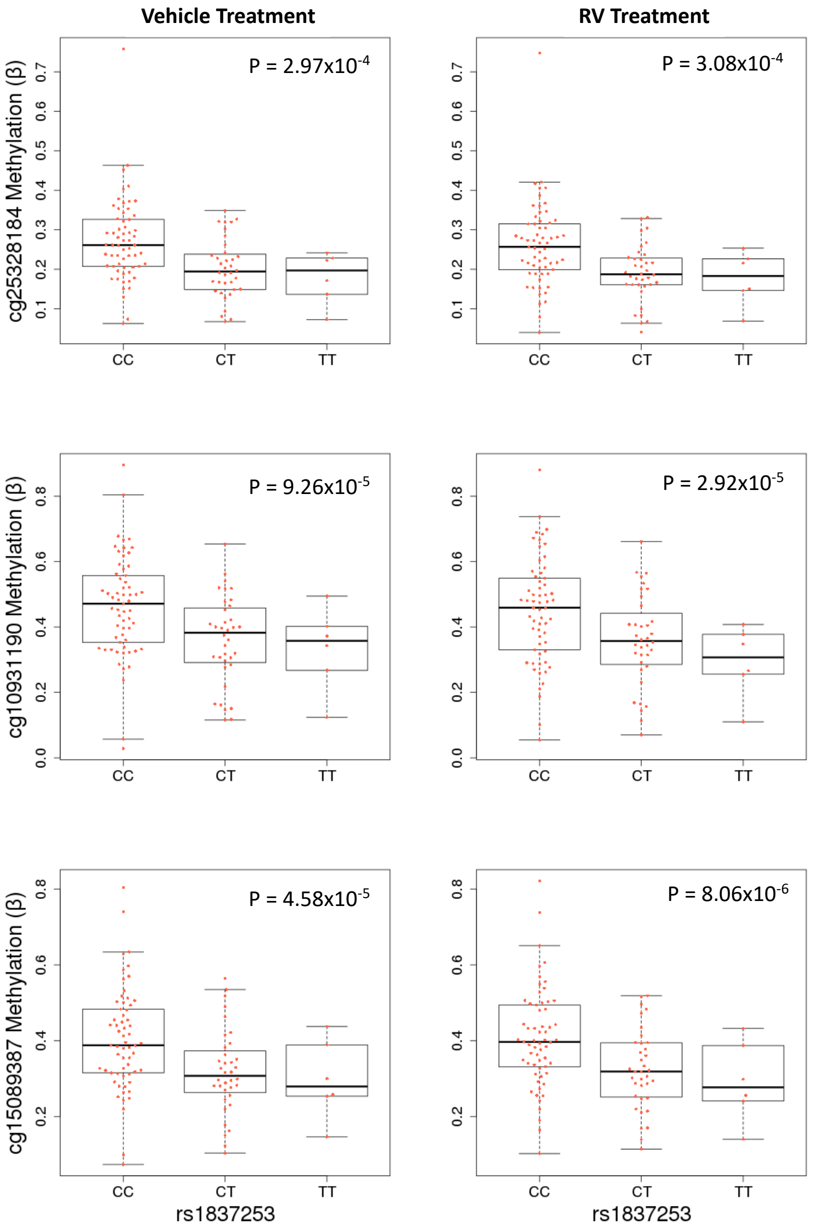
Vehicle Treatment
RV Treatment
cg25328184 Methylation (β)
P = 2.97x10-4
P = 3.08x10-4
cg10931190 Methylation (β)
P = 9.26x10-5
P = 2.92x10-5
cg15089387 Methylation (β)
P = 4.58x10-5
P = 8.06x10-6
CC
CT
TT
rs1837253
